# Fecal Microbial and Metabolic Signatures in VEO-IBD: Implications for Unique Pathophysiology

**DOI:** 10.1101/2025.04.26.650779

**Authors:** Kine Eide Kvitne, Simone Zuffa, Vincent Charron-Lamoureux, Ipsita Mohanty, Abubaker Patan, Helena Mannochio-Russo, Jasmine Zemlin, Lindsey A. Burnett, Lisa S. Zhang, Mia C. Cecala, Ceylan Ersoz, James A. Connelly, Natasha Halasa, Maribeth Nicholson, Pieter C. Dorrestein, Shirley M. Tsunoda, Janet Markle

## Abstract

**Background and Aims:** Very early onset inflammatory bowel disease (VEO-IBD) is a clinically distinct form of IBD manifesting in children before the age of six years. Disease in these children is especially severe and often refractory to treatment. While previous studies have investigated changes in the fecal microbiome and metabolome in adult and pediatric IBD, insights in VEO-IBD remain limited. This multi-omics analysis reveals changes in the fecal microbiome and metabolome in VEO-IBD compared with healthy controls.

**Methods:** Fecal samples were collected from children diagnosed with VEO-IBD and age- and sex-matched healthy controls. Both the fecal metabolome and microbiome were profiled in each sample, using untargeted liquid chromatography coupled with tandem mass spectrometry (LC-MS/MS) and 16S rRNA gene amplicon sequencing.

**Results:** Fecal microbial and metabolic profiles in VEO-IBD were significantly different from healthy controls. Untargeted metabolomics analysis identified a depletion of short-chain *N*-acyl lipids and an enrichment of dipeptides, tripeptides, and oxo bile acids in VEO-IBD patients. Differential abundance analysis of the gut microbiome showed lower abundance of beneficial bacteria such as *Bifidobacterium* and *Blautia*, and higher abundance of *Lachnospira, Veillonella*, and *Bacteroides* in VEO-IBD. The joint analysis suggested a clear association between the altered gut microbiome composition and metabolic dysregulation, specifically for the *N*-acyl lipids.

**Conclusions:** This study offers unique insight into fecal microbial and metabolic signatures in VEO-IBD, paving the way for a better understanding of disease patterns and thereby more effective treatment strategies.

## INTRODUCTION

Inflammatory bowel disease (IBD) comprises a group of chronic conditions characterized by inflammation of the gastrointestinal (GI) tract, with ulcerative colitis (UC) and Crohn’s disease (CD) as the two main subtypes^1^. Very early onset inflammatory bowel disease (VEO-IBD) is a form of IBD manifesting in children younger than six years of age^2,3^. Compared to IBD diagnosed at later ages, VEO-IBD is generally associated with a more severe disease course^4,5^, delayed time to accurate diagnosis^6,7^, and lower initial response rate to conventional treatments^8^. The prolonged exposure to chronic inflammation is of significant concern for key developmental processes such as growth, bone health, and immune function in children with VEO-IBD^9^. Additionally, monogenic etiologies— mutations in a single gene— play a larger role in the pathogenesis of VEO-IBD, especially in infantile onset cases (<2 years)^7,10^. The incidence of VEO-IBD is rapidly increasing worldwide^11,12^, with the majority of children with VEO-IBD (70-80%) not having an identified genetic etiology^13^. This underscores the need to better understand its multifactorial etiology, driven by complex interactions between host, microbial, and environmental factors^14,15^.

The gut microbiome— a key modulator of many physiological processes, including host digestion and vitamin synthesis^16^, drug metabolism^17^, immune function^18^, and metabolic homeostasis^19^— is increasingly recognized for its important role in the pathophysiology of IBD. Its role may be particularly relevant in children with VEO-IBD, as the gut microbiome is immature, less diverse, and undergoes rapid changes during the first years of life^20^. It was recently demonstrated that there is a lower abundance of beneficial bacteria belonging to *Bifidobacterium, Collinsella*, and *Akkermansia*, and a higher abundance of potential pathogens such as *Escherichia, Ruminococcus, Clostridium*, and *Veillonella* in children with VEO-IBD compared to age-matched healthy controls^21^. Alterations in the gut microbial profiles can disrupt the production and modulation of microbial metabolites that act as signaling molecules, thereby influencing key processes such as immune function and homeostasis. Indeed, the levels of several metabolites belonging to the classes of bile acids (BAs)^22,23^, short-chain fatty acids (SCFA)^24,25^, sphingolipids^26^, and amino acids (AA)^25,26^ have been shown to be altered in IBD.

To the best of our knowledge, no multi-omics study has been previously conducted in VEO-IBD, limiting our understanding of how the gut microbiome and metabolome may contribute to this disease. In this study, we therefore combined untargeted metabolomics and 16S rRNA gene amplicon sequencing to investigate characteristics of the fecal metabolome and microbiome in children with VEO-IBD compared with healthy age- and sex-matched controls. We demonstrate an altered gut microbiome composition in VEO-IBD, combined with altered levels of important signaling molecules belonging to the classes of *N*-acyl lipids, small peptides, and BAs. We also show, through a joint analysis, a link between the changes in microbiome composition and metabolic alterations in VEO-IBD. These findings will help advance our understanding of the distinct biology of VEO-IBD and support the development of targeted diagnostic and therapeutic strategies.

## MATERIALS AND METHODS

### Participants

The study recruited 28 children initially diagnosed with isolated VEO-IBD and followed at Monroe Carell Jr. Children’s Hospital at Vanderbilt in the Pediatric Gastroenterology, Hepatology, and Nutrition clinic, and 28 healthy age- and sex-matched controls. However, two individuals in the VEO-IBD group were subsequently excluded due to their clinical phenotype, i.e. disease evolved to systemic autoimmunity and/or autoinflammation. The remaining 26 VEO-IBD patient samples were included in subsequent analyses. The healthy controls were not immunocompromised and were free of any parental-reported vomiting or diarrhea for at least 14 days prior to providing a stool specimen. Exact age at the time of sample collection was not available for the healthy controls, however each control was matched to a VEO-IBD patient of similar age (±1 year). All participants were recruited at Vanderbilt University Medical Center (VUMC).

### Ethics approval and informed consent

The study was approved by the Institutional Review Board at VUMC (studies #170067, #202017, #200412, and #111296) and performed in accordance with the ethical standards of the institution and with the Declaration of Helsinki. Written informed consent was obtained from all parents/legal guardians and assent of pediatric participants prior to any study procedures.

### Fecal specimen collection

Fecal material from each child with VEO-IBD was collected at home by the participants’ parents following a single bowel movement. Parents received oral and written instructions on how to collect fecal material into a sterile plastic container, which was then stored in a home freezer until transport to VUMC. For healthy controls, stool samples were collected either at VUMC during the visit, or if not possible, a courier was sent to the home to collect the sample. Each sample was transported to VUMC within 24 hours of collection, aliquoted into 1.5 mL cryovials, and stored at -80 ° until processing.

### Untargeted metabolomics sample preparation

50% methanol (MeOH:H_2_O) was used to extract metabolites from the fecal samples. Briefly, 800 µL of cold 50% MeOH was added to each sample, before they were homogenized at 25 Hz for 5 min on a tissue lyzer, incubated for 30 min at 4 °C, and centrifuged at max speed (15,000 x g) for 10 min at 4 °C. Supernatant (440 µL) was then collected, dried overnight in a CentriVap, and stored at -80 °C for later acquisition. Prior to instrumental analysis, the samples were reconstituted in 200 µL 50% methanol with 1 μM sulfadimethoxine as the internal standard.

### Untargeted metabolomics data acquisition

Samples (5 μL) were injected into a Vanquish ultra-high-performance liquid chromatography (UHPLC) system coupled to a QExactive quadrupole orbitrap (Thermo Scientific) mass spectrometer. The chromatographic separation was done on a polar C18 column (Kinetex Polar C18, 100 mm x 2.1 mm, 2.6 µm particle size, 100 A pore size; Phenomenex) with a matching Guard cartridge (2.1 mm) at 40 °C column temperature. The mobile phase consisted of solvent A H_2_O + 0.1% formic acid (FA), and solvent B acetonitrile (ACN) + 0.1% FA. Samples were eluted at a flow rate of 0.5 mL/min using the following gradient: 0–0.5 min, 5% B, 0.5–1.1 min, 5–25% B, 1.1–7.5 min, 25-60% B, 7.5–8.5 min, 60-99% B, 8.5–9.5 min, 99% B, 9.5–10 min, 99-5% B, 10.0–10.5 min 5% B, 10.5–10.75, 5-99% B, 10.75–11.25 min, 99% B, 11.25–11.5 min, 99-5% B, 11.5–12 min, 5% B. All solvents used were LC-MS grade. Data dependent acquisition (DDA) of tandem mass spectrometry (MS/MS) spectra was performed in positive mode with the following parameters: sheath gas flow 53 L/min, aux gas flow rate 14 L/min, sweep gas flow 3 L/min, spray voltage 3.5 kV, inlet capillary 269 °C, aux gas heater 438 °C, and 50 V S-lens level. The MS scan range was set to 150–1000 *m/z* with a resolution of 35,000 at *m/z* 200; maximum ion injection time, 100 ms; automatic gain control (AGC) target, 5.0E4. MS/MS spectra were collected with a resolution of 17,500 and an AGC target of 5E5 with a maximum injection time of 150 ms, and fragmented the top 5 most abundant ions per cycle with a 1 *m/z* isolation window, isolation offset set to 0 *m/z*, stepped collision energies of 25, 40, 60, and a dynamic exclusion window of 10 s.

### Untargeted metabolomics data processing

The acquired .raw files were converted into .mzML format using MSConvert^27^. Feature detection and extraction were performed using MZmine 4.2^28^. For mass detection, MS1 and MS2 noise levels were set to 5.0E4 and 1.0E3, respectively. Parameters for the chromatogram builder were set to minimum 5 consecutive scans, 1.0E5 minimum absolute height, and *m/z* tolerance 10 ppm. Parameters for the local minimum resolver function were set to 85% for the chromatographic threshold, 0.2 min for minimum search range RT, and 1.7 for minimum ratio of peak top/edge. 13C isotope filter and finder were applied. Join aligner was used to align features with weight for *m/z* set to 80 and RT tolerance set to 0.2 min. Before using the peak finder function, features not detected in at least 2 samples were removed. metaCorrelate and ion identity networking were performed. The .mgf spectra file and associated feature table were then exported and used to generate a feature based molecular network (FBMN)^29^ on GNPS2^30^. Briefly, fragment tolerances for both parent and fragment ions were set at 0.02, while networking and annotation parameters were set to minimum 5 matching peaks and cosine similarity > 0.7. The GNPS2 FBMN job is available at https://gnps2.org/status?task=8f1989e83449460a9a5748cd8f32df30. The network.graphml file from the FBMN was imported and manipulated in Cytoscape^31^ (version 3.10.3). MS/MS spectra of unknown features were imported into SIRIUS^32^ (version 6.0.7) and CANOPUS^33^ was used to predict their chemical pathways or classes. Pathways or classes were retained if probability scores were > 0.7. Known and unknown features of interest were also searched using two domain-specific MASSTs^34^, microbeMASST^35^ (https://masst.gnps2.org/microbemasst/) and tissueMASST (https://masst.gnps2.org/tissuemasst/). MicrobeMASST is a taxonomically informed mass spectrometry search tool within the GNPS ecosystem where MS/MS spectra can be linked to their respective microbial producers by querying against a curated reference database of >60,000 microbial monocultures. TissueMASST, a similar MS/MS search tool, enables investigation of molecules of interest across diverse biological contexts by the search of MS/MS spectra across publicly available metabolomics datasets acquired from preclinical animal models (mouse and rat) and humans and with associated metadata on tissue localization and disease status.

### Untargeted metabolomics data analysis

Metabolomics data was then imported in R 4.4.2 (R Foundation for Statistical Computing, Vienna, Austria) for downstream data analysis. Quality control (QC) samples were used to evaluate data quality. Features with a retention time (RT) < 0.70 or > 9.5 min were excluded from the analysis. Because polymers were detected in the LC-MS/MS run, the package `homologueDiscoverer v 0.0.0.9000`^36^ was used to remove them. Six samples (VEO-IBD; n = 5, healthy; n = 1) were excluded from the metabolomics analysis because of low intensity in the total ion chromatogram (TIC; n = 3) or high contamination (n = 3). Blank subtraction was performed by removing features detected in blank samples where the mean peak areas were less than five times that observed in the pooled QC samples. Features with near zero variance were removed using the package `caret v 6.0`. A validation of features annotated as BA candidates was performed using massQL^37–39^. Multivariate analysis was conducted using the package `mixOmics v 6.30`. Principal component analysis (PCA) and partial least square discriminant analysis (PLS-DA) were performed on peak areas after robust center log ratio transformation (rclr) using the package `vegan v 2.6.10`. PERMANOVA was used to evaluate group centroid separation. The performance of the PLS-DA models were evaluated using a 4-fold cross-validation. Variable importance in projection (VIP) scores were calculated per feature and features with VIP > 1 were considered significant. Natural log ratios of features of interest were generated by summing the peak areas. Wilcoxon rank sum test, followed by Benjamini-Hochberg method to adjust for multiple comparisons, was used to investigate differences between the VEO-IBD group and the control group. Chi-squared test was used to compare categorical variables between the two groups. P < 0.05 was considered statistically significant. Pearson correlation was used to investigate correlation between age and alpha diversity.

### 16S rRNA gene amplicon sequencing and processing

The UC San Diego Microbiome Core performed DNA extraction and sequencing following the previously developed standard protocols from the Earth Microbiome Project (http://earthmicrobiome.org/protocols-and-standards/16s/)^40^. Briefly, extraction was conducted using the MagMAX Microbiome Ultra Nucleic Acid Isolation Kit (Thermo Fisher Scientific, USA) and KingFisher Flex robots (Thermo Fisher Scientific, USA). Negative (blanks) and positive (Zymo mock communities) controls were included in the analysis and used for quality controls. Sequencing was conducted on the V4 region of 16S rRNA gene using the 515F forward and the 806R reverse primers on a MiSeq System (Illumina, USA). Generated demultiplexed fastq files were then imported in Qiita^41^ (ID 15748) for processing. Briefly, the Qiita default workflow for 16S rRNA data was applied, which included trimming to 150 base pairs and Deblur for the generation of the final OTU (operational taxonomic unit) table. Taxonomic classification was also performed in Qiita using the Greengenes 13.8 phylogeny database^42^. The obtained .biom file was then converted into a .tsv table using QIIME 2^43^ and imported in R 4.4.2 for downstream analysis.

### Microbiome data analysis

The sequencing depth of the microbiome data was inspected and five samples (VEO-IBD; n = 1, healthy; n = 4) were excluded from the downstream analysis due to low read counts (< 9,000 reads). The `phyloseq v 1.50`^44^ package was used to manipulate the microbiome data. Alpha diversity analysis was conducted by rarefying the data to the lowest number of reads observed in the cohort and calculated using the Shannon Diversity Index. Before ordination and differential abundance analysis, OTUs were collapsed at genus level. Beta diversity analysis was conducted via PCA of the rclr transformed data. Significant differences in community profiles were tested via PERMANOVA^45^. A PLS-DA model was also generated to extract features driving group separation. The performance was evaluated using a 4-fold cross-validation, and features with VIP > 1 were considered significant. Finally, differential abundance analysis was performed using ALDEx2^46^ and features with adjusted p value < 0.05 were considered significant.

### Multi-omics analysis

The joint analysis between the microbiome and the metabolomics data was performed using DIABLO^47^, a supervised method using multi-block PLS-DA to identify feature correlations between datasets in relation to a categorical outcome. The model was first tuned to retain top features for each omics block maximizing discriminatory covariance between groups. Model performance was evaluated via leave-one-out cross validation. A circos plot was finally generated to display intra feature correlation with a cut off set to 0.5.

## RESULTS

A total of 26 children previously diagnosed with VEO-IBD and 28 healthy age- and sex-matched controls were included in this study. One stool sample per subject was collected and used for 16S rRNA gene amplicon sequencing, to profile the fecal microbial communities, and untargeted liquid chromatography coupled with tandem mass spectrometry (LC-MS/MS) to profile both known and unknown metabolites (**Fig. 1a**). Mean age ± SD of study participants were 6.7 ± 3.4 and 6.4 ± 3.5 years in the VEO-IBD group and control group (Wilcoxon test: p = 0.71), respectively, with 54% being male in both groups (Chi-squared test: p =1). Demographic characteristics of the study participants are listed in **Table 1**. Most patients with VEO-IBD (85%) received pharmacological treatment, including tumor necrosis factor alpha (TNF-α)-inhibitors (54%), aminosalicylates (31%), antibiotics (8%), or a combination of these therapies; and 4 subjects were diagnosed with monogenic VEO-IBD. Mean time from VEO-IBD diagnosis to sample collection was 3.3 ± 3.2 years, with 46% of samples being collected within the first year.

**Table 1.** Characteristics of the study population.

| Characteristics | VEO-IBD<br>(n = 26) | Controls<br>(n = 28) |
| --- | --- | --- |
| Age at enrollment (years) | 6.7 (1-14) | 6.4 (1-14) |
| Sex (female/male) | 12/14 | 13/15 |
| Monogenic IBD, n (%) | 4 (15%) | NA |
| Pharmacological treatment, n (%) | 22 (85%) | NA |
| - Aminosalicylate (sulfasalazine, 5-ASA, balsalazide) | 8 (31%) |  |
| - TNF-α-inhibitor (infliximab, adalimumab) | 14 (54%) | |
| - Antibiotics (amoxicillin, azithromycin) | 2 (8%) |  |

Numbers are given as mean (absolute range) or count (%). Abbreviations: 5-ASA, 5-aminosalicylate; NA, not applicable; TNF-α , tumor necrosis factor alpha

**Figure 1.**
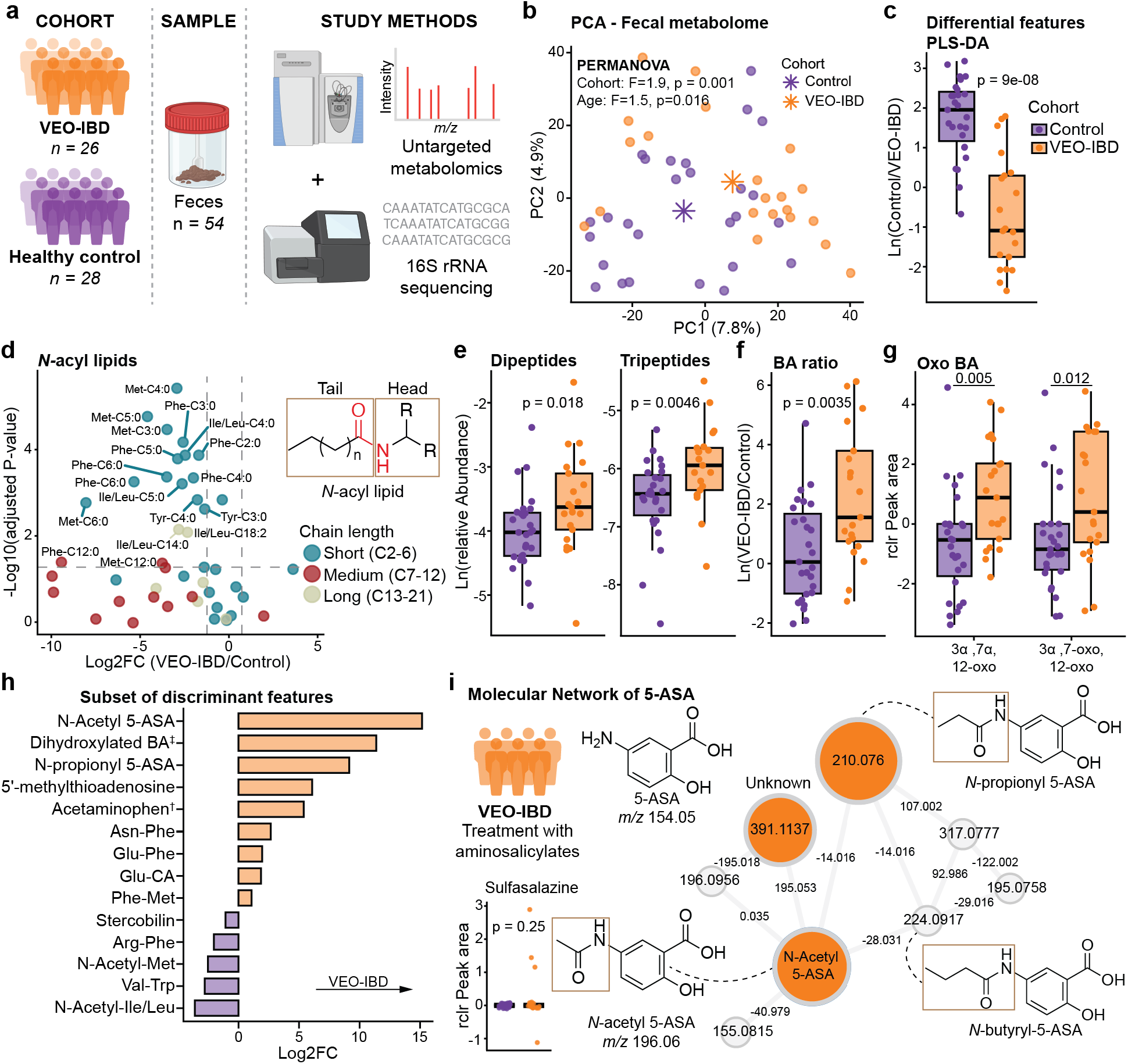
*N-*acyl lipids, small peptides, oxo bile acids, and drug metabolites of aminosalicylates are main drivers of separation in the fecal metabolome between VEO-IBD and controls. Overview of the study design. Twenty six patients with VEO-IBD and 28 age- and sex-matched healthy controls provided one stool sample for untargeted LC-MS/MS analysis and 16S rRNA gene amplicon sequencing. **(b)** PCA of fecal metabolic profiles showed separation based on cohort (PERMANOVA, R^2^ = 0.039, F-statistic = 1.9, p = 0.001) and age (PERMANOVA, R^2^ = 0.031, F-statistic = 1.5, p = 0.016). **(c)** A PLS-DA model was constructed to extract features separating VEO-IBD and control. Boxplot highlights the natural log ratio of differential features from the PLS-DA model (numerator = control, denominator = VEO-IBD). **(d)** Volcano plot of *N-*acyl lipids, with the dotted lines showing the significance threshold (p adjusted (Benjamini-Hochberg) < 0.05 and absolute log2FC > 2). **(e)** Boxplot of the relative abundance of dipeptides and tripeptides with a VIP score > 1 from the PLS-DA model, respectively. Relative abundance was calculated by dividing the sum of peak areas for relevant dipeptides and tripeptides by the total ion current. **(f)** Boxplot of the natural log ratio of BAs with a VIP score > 1 from the PLS-DA model (numerator = VEO-IBD, denominator = control). **(g)** Boxplot of the annotated oxo Bas - 3α-7α-12-oxo (Wilcoxon test, p = 0.005) and 3α-7-oxo-12-oxo (Wilcoxon test, p = 0.012). P values were adjusted for multiple comparisons using the Benjamini-Hochberg method. **(h)** Barplot of other differential features between VEO-IBD and control (VIP score > 1). **(i)** Molecular network of 5-ASA, the major metabolite of sulfasalazine and balsalazide, and its drug metabolites. Orange nodes represent features with a VIP > 2, with larger nodes indicating higher VIP scores. All boxplots show the first (lower), median, and third (upper) quartiles, with whiskers 1.5 times the interquartile range. ^‡^Candidate dihydroxylated BA (Δ 135.033). ^†^MS/MS spectral match to 2-acetamidophenol, a stereoisomer of acetaminophen, which is indistinguishable from acetaminophen via mass spectrometry.

### Children with VEO-IBD display distinct metabolic profiles

Using untargeted LC-MS/MS metabolomics, we observed a significant separation in the fecal metabolome between the VEO-IBD group and the controls (**Fig. 1b**) (PERMANOVA: R^2^ = 0.039, F-statistic = 1.9, p = 0.001). Age also had a significant impact on the fecal metabolomic profiles (PERMANOVA: R^2^ = 0.031, F-statistic = 1.5, p = 0.016). To identify discriminant metabolites between the two groups, a supervised Partial Least Squares - Discriminant Analysis (PLS-DA) model was generated (**SI Fig. 1a**, classification error rate (CER) = 0.21) and 1,678 known and unknown metabolic features with variable importance in projection (VIP) > 1 were extracted (**SI Table 1**). The natural log ratio of the extracted features significantly separated the controls from the VEO-IBD group (Wilcoxon test, p = 9.0e-08) (**Fig. 1c**). One group of metabolites that stood out as key discriminant features between VEO-IBD and controls were the *N-*acyl lipids, signaling molecules containing an amine group (head) and a fatty acid (tail), linked by an amide bond^38^ (**Fig. 1d**). Several putatively annotated short-chain (C2-C6) *N-*acyl lipids with methionine (Met), phenylalanine (Phe), leucine (Leu)/isoleucine (Ile), and tyrosine (Tyr) as head groups were significantly depleted in children with VEO-IBD (**Fig. 1d, SI Fig. 1b**). In contrast, we observed a higher relative abundance of dipeptides and tripeptides in VEO-IBD than in controls (Wilcoxon test: p = 0.018 and p = 0.0046, respectively) (**Fig. 1e**). Annotated dipeptides that were enriched in VEO-IBD included asparagine (Asn)-Phe, glutamic acid (Glu)-Phe, Phe-Met, Ile-Leu, Trp-Phe, Phe-Leu, Tyr-Phe, valine (Val)-Ile, threonine (Thr)-Phe, Met-Leu, Ile-Met, aspartic acid (Asp)-Phe, and Ala-Phe (**SI Table 1**). Annotated tripeptides included Ile-Proline(Pro)-Ile and Thr-Val-Leu. Given the limited number of annotated tripeptides in our dataset, we also included unknown features predicted to be tripeptides by SIRIUS (probability > 0.7) in the analysis. A total of 43 tripeptides with a VIP > 1 were identified, and a group comparison of the relative abundance supported higher levels of tripeptides in VEO-IBD (Wilcoxon test: p = 0.00011, **SI Fig. 1c**).

Several annotated and putatively annotated BAs were important discriminant features (Wilcoxon test, p = 0.0035) (**Fig. 1f**). Notably, children with VEO-IBD displayed a higher abundance of oxo BAs compared with controls, specifically putative 3α-7α-12-oxo and 3α-7-oxo-12-oxo (**Fig. 1g**). Among the other annotated oxo BAs, there was a trend towards a higher abundance of three different 3α-7α-12-oxo isomers and 3α-7-oxo-12-oxo in VEO-IBD, while 3α-12-oxo did not differ between the groups (**SI Fig. 1d**). Microbial AA conjugated BAs such as Glu-cholic acid (CA) was also enriched in VEO-IBD (**Fig. 1h**). We did not observe any difference in host-derived CA, microbially derived deoxycholic acid (DCA), or taurine or glycine conjugated BAs (**SI Fig. 1e**).

Additional discriminant metabolic features enriched in VEO-IBD included drug metabolites of the aminosalicylates, acetaminophen/2-acetamidophenol (stereoisomers and thereby indistinguishable via mass spectrometry), a candidate dihydroxylated BA (refers to a BA with uncharacterized modification)^37^, and 5’-methylthioadenosine. On the other hand, *N*-acetyl-Met, *N*-acetyl-Ile/Leu, Arg-Phe, Val-Trp, and stercobilin were enriched in control (**Fig. 1h**). It should be noted that we observed poor alignment (38%) between acetaminophen detection and reported use, possibly due to limited MS/MS fragmentation affecting annotation or inaccuracies in self-reported use of acetaminophen. Around 30% of the children with VEO-IBD reported use of an aminosalicylate. Using the GNPS drug library^48^, a resource encompassing MS/MS data of drugs and their metabolites/analogs, combined with a molecular network of the aminosalicylates, we were able to annotate *N*-acetyl 5-ASA and *N*-propionyl 5-ASA as key discriminant features between VEO-IBD and controls (**Fig. 1i**). These gut microbially derived metabolites^49^ were detected in all patients reported to receive treatment with an aminosalicylate. 5-aminosalicylate (5-ASA) eluted in the dead volume. Sulfasalazine was not a discriminant feature (Wilcoxon test: p = 0.25) and no spectral matches to balsalazide were observed, highlighting the importance of also considering drug metabolites. Several other unannotated metabolic features were also driving the separation between VEO-IBD and controls. To predict these metabolites’ chemical classes, their MS/MS spectra were processed with SIRIUS. Features with a VIP > 2.5 were further investigated. Of the metabolites enriched in VEO-IBD, many were predicted to be dipeptides and tripeptides, while depleted metabolites included piperidine, purine, and simple indole alkaloids (**SI Fig. 2a**). To investigate their possible microbial origin, we also searched the MS/MS spectra via microbeMASST (see methods). The output showed that out of the 20 most discriminant features (top 10 in each group), 14 have been previously observed in microbial monocultures.

### Gut microbiome composition is altered in VEO-IBD versus controls

We next looked at the microbial alpha diversity by measuring the Shannon Diversity Index, and found no statistical significant difference between VEO-IBD and controls (Wilcoxon test, p = 0.48) (**Fig. 2a**). However, when investigating age, we observed a moderate positive correlation between age and the Shannon Diversity Index in the VEO-IBD group (Pearson, R = 0.41, p = 0.041), but not in the control group (p = 0.33) (**Fig. 2b**). There was also a significant dissimilarity in beta diversity based on PCA of the robust center log ratio (rclr) transformed data (PERMANOVA, R^2^ = 0.087, F-statistic = 4.5, p < 0.001) (**Fig. 2c**). A PLS-DA model was generated to identify discriminant taxa at genus level (**SI Fig. 3a**, CER = 0.11). We identified 32 genera with VIP > 1; 17 that were enriched in the control group and 15 that were enriched in the VEO-IBD group (**SI Table 2**). Natural log ratios of the selected taxa confirmed a significant difference in the fecal microbiome composition between the two groups (Wilcoxon test, p = 3.1e-09) (**Fig. 2d**). Key discriminant genera, with a higher abundance in VEO-IBD, included *Lachnospira, Sutterella, Clostridium, Bacteroides, Alistipes*, and unknown genera of the *Enterobacteriaceae* family (**Fig. 2e, SI Fig. 3b**). On the other hand, genera including *Coprococcus, Bifidobacterium, Collinsella, Blautia*, and *Dorea* were depleted in VEO-IBD. To further investigate differences in fecal microbiome composition, differential abundance analysis was also conducted using ALDEx2^46^. The output showed similar findings as from the PLS-DA model, with higher abundance of *Veillonella, Sutterella, Lachnospira, Clostridium*, and *Bacteroides* and lower abundance of *Coprococcus, Collinsella*, and *Turibacter* in VEO-IBD compared with controls (**Fig. 2f**).

**Figure 2.**
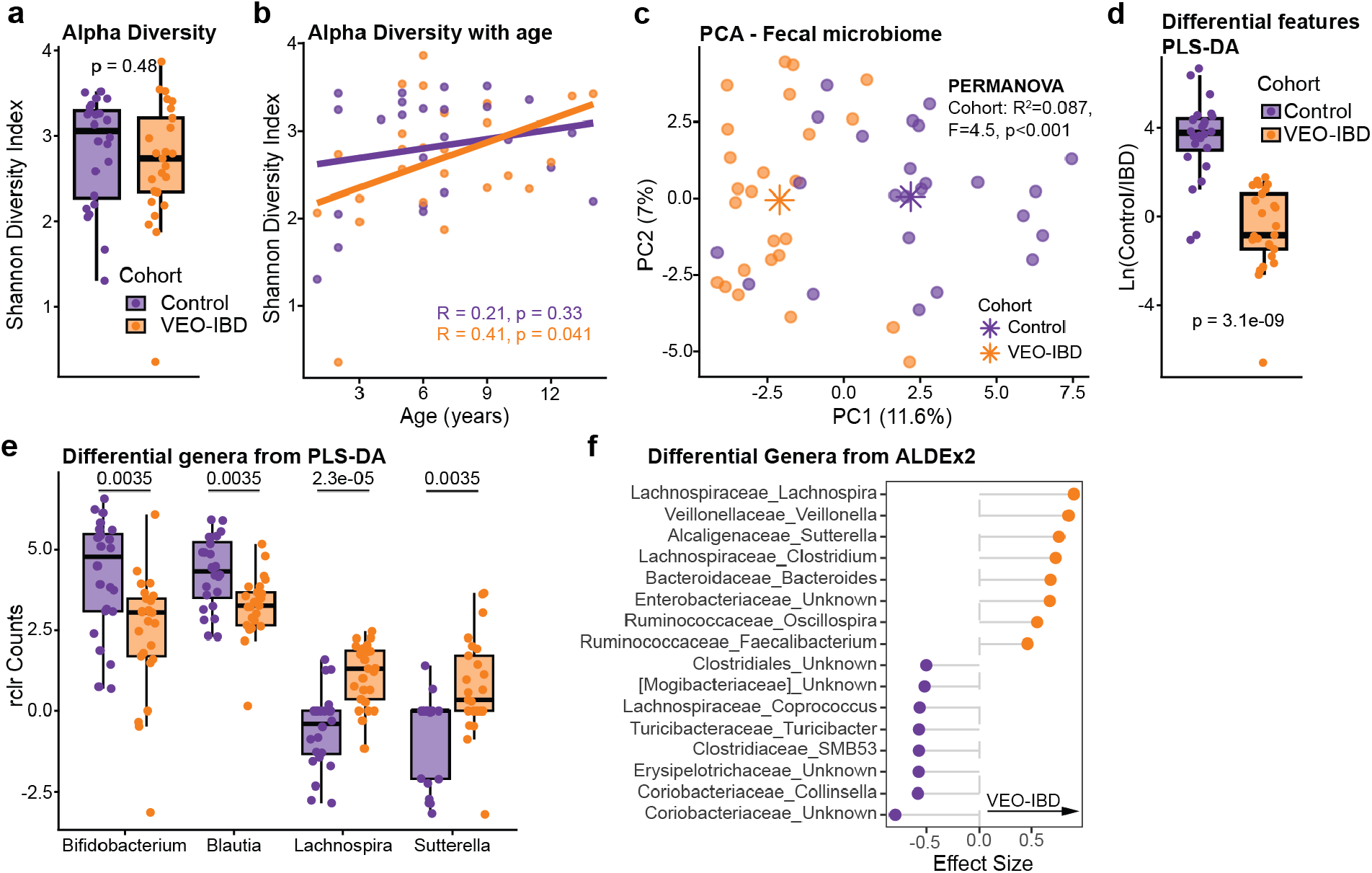
Fecal microbiome composition varies significantly between VEO-IBD and controls. **(a)** Boxplot of alpha diversity measured as Shannon’s Diversity Index (Wilcoxon test, p = 0.48). **(b)** Scatter plot showing the correlation between age and Shannon’s Diversity Index (Pearson’s correlation, VEO-IBD: R = 0.41, p = 0.041; control: R = 0.21, p = 0.33). **(c)** PCA of rclr transformed data showing a significant difference in beta diversity between the two cohorts (PERMANOVA, R^2^ = 0.087, F-statistic = 4.5, p < 0.001) with no effect of age. **(d)** Boxplot of the natural log ratio of differential genera extracted from the PLS-DA model (numerator = control, denominator = VEO-IBD). **(e)** Boxplot of selected differential genera obtained from the PLS-DA model. P values were adjusted for multiple comparisons using the Benjamini-Hochberg method. **(f)** Balloon plot of differential abundance taxa collapsed at genus level using ALDEx2. Only taxa with adjusted p value < 0.05 are reported. Taxa names reported as Family_Genera. All boxplots show the first (lower), median, and third (upper) quartiles, with whiskers 1.5 times the interquartile range.

### Joint analysis identifies link between *N-*acyl lipid dysregulation and the gut microbiome

To investigate if any of the metabolic changes we detected could be explained by changes in the composition of the gut microbiome, a multi-omics analysis was performed using DIABLO^47^. The output of the first component (correlation cut-off > 0.5) showed a correlation between the short-chain *N-*acyl lipids Met-C3:0, Met-C4:0, Phe-C3:0, Phe-C4:0, and Ile/Leu-C4:0, and the genera *Coprococcus, Lachnospira, Collinsella, Bifidobacterium, Blautia, Dorea, Alistipes*, and *Bacteroides* (**Fig. 3a**). More specifically, *N-*acyl lipids were positively correlated with *Coprococcus, Collinsella, Bifidobacterium, Blautia*, and *Dorea*, genera more abundant in the healthy controls, and negatively correlated with *Lachnospira, Bacteroides*, and *Alistipes*, enriched in VEO-IBD. Similarly, *N*-acetyl-Ile/Leu and *N*-acetyl-Met, both more abundant in controls, were positively correlated with *Coprococcus, Collinsella, and Blautia* and inversely correlated with *Lachnospira* and *Alistipes*. Additionally, we observed correlations between specific genera and unannotated metabolic features (**Fig. 3a**). Constructed sub-molecular networks did not contain any annotated features. SIRIUS predictions indicated that these unknown metabolic features belonged to the pathways or classes of alkaloids (*m/z* 243.1128, *m/z* 197.1284, and *m/z* 188.128), dipeptides (*m/z* 395.1811), and tripeptides (*m/z* 346.1975) (**Fig. 3b**). Next, we searched them via microbeMASST and found that three of them have been previously observed in microbial monocultures. Of particular interest was the unknown feature with *m/z* 346.1975 enriched in VEO-IBD, which SIRIUS-predicted to be a tripeptide, matching to genera such as *Bacteroides* and *Akkermansia* (**SI Fig. 3c**). We also searched this feature (*m/z* of 346.1975) via tissueMASST to investigate its presence in other publicly available metabolomics IBD datasets. The output showed that this metabolic feature is detected with higher frequency in feces of individuals with IBD compared with healthy (Chi-squared test: p = 2.2e-16), and also in other chronic inflammatory diseases such as rheumatoid arthritis compared with healthy (Chi-squared test: p = 5.106e-08) (**Fig. 3b**).

**Figure 3.**
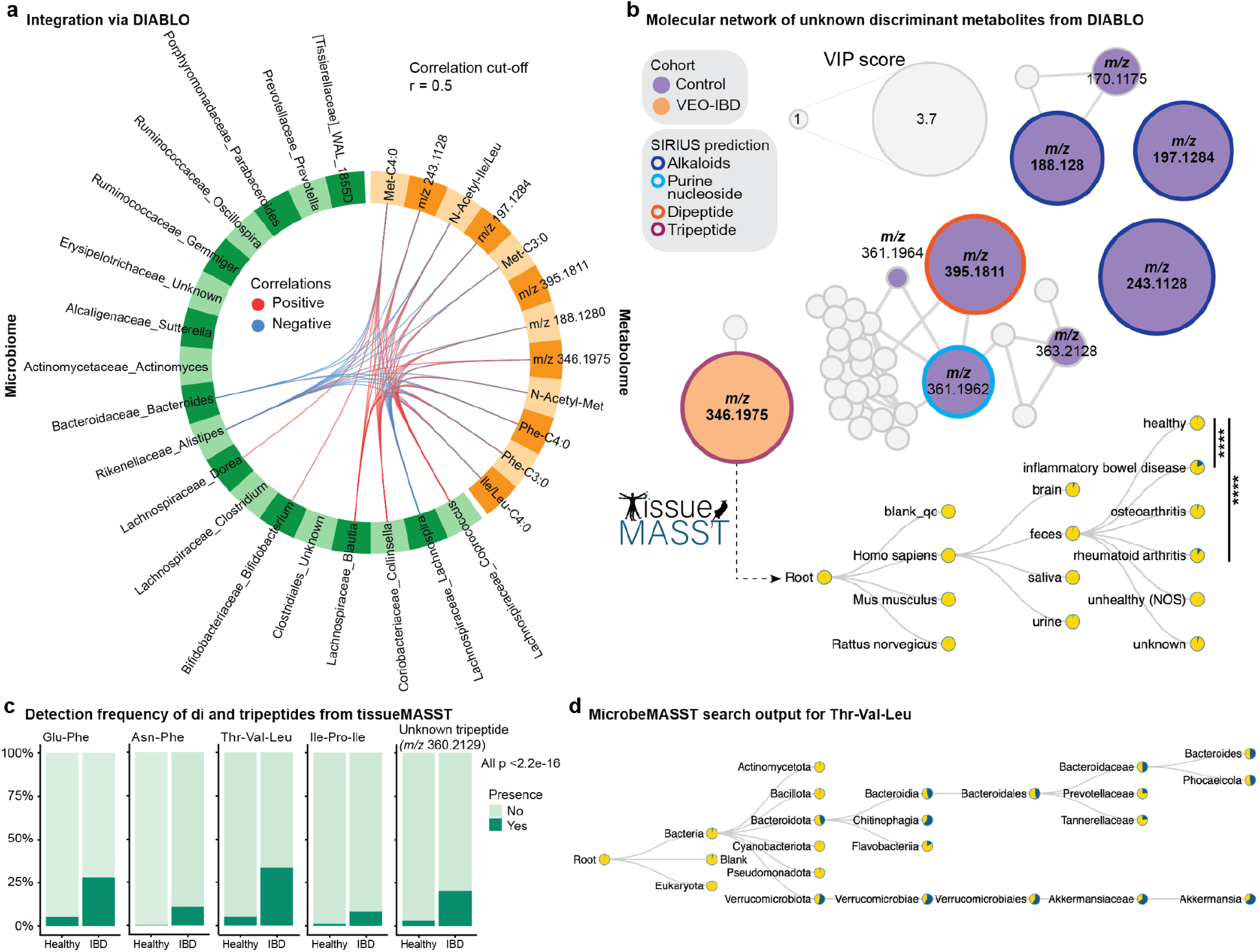
Multi-omics analysis of the microbiome and metabolome data reveals significant correlation between specific microbial genera, *N*-acyl lipids, and unknown metabolic features. **(a)** Integration of the metabolomics and microbiome data using DIABLO. The correlation cut-off was set to > 0.5. Red lines show positive correlation, blue lines show negative correlation. **(b)** Sub-molecular networks of unknown metabolic features highlighted by DIABLO. MS/MS spectra of the unknown metabolites were processed using SIRIUS to predict their chemical pathways or classes. The MS/MS spectra of one of the unknown features enriched in VEO-IBD (*m/z* 346.1975), predicted to be a tripeptide, was searched across publicly available metabolomics datasets using tissueMASST. The output showed a significantly higher detection rate of this feature in fecal samples from individuals with IBD compared to healthy (Chi-squared test: p < 2.2e-16). Pie charts show the proportion of spectral matches found in deposited datasets, with blue representing a match and yellow representing a non-match. **(c)** Other key discriminant features belonging to the classes of di- and tripeptides were therefore also searched via tissueMASST. Barplots showing a higher detection frequency in human fecal samples from individuals with IBD than healthy individuals (Chi-squared test: all p < 2.2e-16). The Y-axis shows the percentage of samples where the feature was detected vs not detected. **(d)** To investigate if they are of microbial origin, these features were also searched against microbeMASST. The output for Thr-Val-Leu is illustrated, but they all matched to microbial monocultures. Pie charts show the proportion of spectral matches found in deposited datasets, with blue representing a match and yellow representing a non-match. Significance: **** p < 0.0001.

This observation motivated us to explore the detection frequency of other discriminant di- and tripeptides from the metabolomics PLS-DA model in other publicly available metabolomics IBD datasets. The MS/MS spectra of Glu-Phe, Asn-Phe, Thr-Val-Leu, Ile-Pro-Ile, and an unknown feature SIRIUS-predicted to be a tripeptide, were therefore searched via tissueMASST. We focused on the output from datasets comprising human stool samples. The output confirmed a higher detection rate of all these features in human fecal samples from individuals with IBD compared with healthy individuals (Chi-squared test: all p < 2.2e-16) (**Fig. 3c**). We also searched the MS/MS spectra of these features via microbeMASST. The output confirmed that they are microbial metabolites. The spectra of these metabolites were most frequently observed in cultures of *Bacteroides, Veillonella*, and *Akkermansia*, although they matched to several bacterial cultures, as illustrated for Thr-Val-Leu (**Fig. 3d**).

## DISCUSSION

A more thorough understanding of gut microbiome composition and function in IBD may set the stage for more effective and personalized treatments. For most patients with IBD, the peak age at onset is between 20-40 years^50^. However, a small but growing percentage of IBD patients are diagnosed with VEO-IBD at age 6 or younger, and disease in these children is especially severe and often treatment-refractory^3^. The timely selection of an individualized treatment plan is critical for these children, since VEO-IBD is associated with poor long-term outcomes, including surgical resection, increased risk of colorectal cancer, and even death^7^. The gut microbiome represents a promising therapeutic target in IBD. Significant research efforts have been dedicated to investigate changes in the gut microbiome and metabolome in pediatric^26,51,52^ and adult IBD^53,54^. However, studies specifically looking at VEO-IBD remain scarce^21^, limiting our understanding of whether gut microbial and metabolic changes may contribute to the distinct clinical phenotype of VEO-IBD. Given that the gut microbiome and immune system evolve rapidly in early childhood^20^, VEO-IBD may involve unique perturbations that differ, not only from healthy peers, but also from those seen in IBD diagnosed at later ages. Untargeted LC-MS/MS metabolomics allows the identification of thousands of known and unknown metabolites in a given sample. When combined with 16S rRNA gene amplicon sequencing, which characterizes the gut microbiome composition, it can provide valuable insights into the intricate relationship between host, microbes, and metabolites. To our knowledge, this is the first multi-omics analysis investigating changes in the fecal microbiome and metabolome in children diagnosed with VEO-IBD compared with healthy age- and sex-matched controls.

*N*-acyl lipids, an understudied class of molecules, appear to play an important role in several biological functions, such as immune regulation^38,55^. In this study, short-chain *N*-acyl lipids were among the top discriminant features between VEO-IBD and healthy participants. Several putatively annotated *N*-acyl lipids had a significantly lower abundance in VEO-IBD compared with controls. This depletion of short-chain *N*-acyl lipids in VEO-IBD compared to healthy controls has, to our knowledge, not been previously described in the literature. Whether this represents a unique characteristic of VEO-IBD or is a shared trait across IBD subtypes is not yet known. Our study also demonstrated a significant correlation between the abundance of these short-chain *N*-acyl lipids and the abundance of specific bacterial genera in a multi-omics analysis of the fecal metabolome and the microbiome. Met-C3:0, Met-C4:0, Phe-C3:0, Phe-C4:0, and Ile/Leu-C4:0 were negatively correlated with bacterial genera enriched in VEO-IBD, and positively correlated with genera more abundant in healthy controls. This pattern suggests that the dysregulation of *N*-acyl lipids in VEO-IBD may be linked to underlying gut microbiome dysbiosis. We recently demonstrated that many of these *N*-acyl lipids are microbially produced when both the amine group and lipid chain are present^38^. We speculate that a lower abundance of specific short-chain fatty acid (SCFA) producing bacteria in VEO-IBD, such as *Coprococcus, Blautia, Dorea*, and *Bifidobacterium*^*56*^, may alter SCFAs availability, thereby limiting their ability to form *N*-acyl lipids with an amine group.

Recent studies have demonstrated a correlation between higher dipeptide abundance and increased disease activity in UC^57^. The elevated dipeptide levels have been linked to an overabundance of microbial proteases – particularly serine proteases and dipeptidases produced by *Bacteroides*, specifically *B. vulgatus*^*57*^. Consistent with this finding, we observed a higher relative abundance of dipeptides and tripeptides combined with a higher abundance of *Bacteroides* in VEO-IBD compared with healthy controls. Furthermore, a microbeMASST search confirmed a higher detection frequency of several di- and tripeptides in cultures of *Bacteroides* and *Veillonella*, both of which were enriched in VEO-IBD. These findings suggest that the elevated levels of small peptides are likely secondary to increased proteolytic activity associated with the altered gut microbiome in VEO-IBD. While it remains unknown whether these peptides directly contribute to VEO-IBD pathophysiology or are simply byproducts of the altered gut microbiome composition, there is evidence pointing to a potential role in disease. For instance, protease inhibitors have been shown to prevent colitis in *B. vulgatus* monocolonized, IL10-deficient mice^57^, suggesting that microbial protease activity could be a viable therapeutic target in IBD. Additionally, a tissueMASST search found that key discriminant di- and tripeptides were more frequently detected in fecal samples from individuals with IBD than from healthy subjects. This suggests the possibility that certain di- or tripeptides may serve as biomarkers for IBD or disease activity and/or severity.

Previous studies have shown dysregulation of BA metabolism in IBD^22,23^. BAs are cholesterol-derived signaling molecules synthesized in the liver that activate receptors such as farnesoid X receptor (FXR) and pregnane X receptor (PXR)^58^. After being conjugated with taurine and glycine, BAs are secreted into the small intestine via the biliary system where they undergo extensive modifications via microbial metabolism^58^. Consistent with previous IBD literature^23,59^, we observed a shift in the bile acid pool in VEO-IBD, characterized by a higher abundance of microbial conjugated BAs such as putative Glu-CA. Bile salt hydrolase (BSH), an enzyme widely expressed across most bacterial phyla^60^, plays a key role in the modification of BAs, including AA conjugation of BA^61,62^. Notably, Glu-CA has been shown to be more resistant to BSH-mediated hydrolysis than other microbial conjugated BAs in an in vitro model with 17 gut bacterial isolates exposed to AA conjugated BAs^63^. We also observed elevated levels of some, but not all, 12-oxo BAs in VEO-IBD. These oxo BAs which contain a ketone in the carbon 12 position in the steroid ring, have been implicated in activating the pyrin inflammasome and promoting secretion of pro-inflammatory cytokines, such as interleukin (IL)-18 and IL-1β^64^. This suggests a potential role for the oxo BAs in immune regulation and disease pathophysiology in VEO-IBD. Interestingly, unlike previous reports in pediatric IBD^51,65^, we did not observe differences in levels of host-derived CA or microbially derived DCA. These findings indicate that BA metabolism in VEO-IBD may follow a distinct pattern compared to IBD diagnosed later in childhood or in adulthood.

In line with previous findings in pediatric IBD^21,66^, we observed significant differences in gut microbiome composition between children with VEO-IBD compared with age- and sex-matched controls. This was evidenced by a significant difference in microbial community dissimilarity between the groups. Specifically, we found a significantly lower abundance of beneficial commensal gut bacteria including *Bifidobacterium, Blautia*, and *Coprococcus* – microorganisms associated with host health^67,68^. Supporting this, Conrad et al. recently reported increased microbial dissimilarity in VEO-IBD compared with IBD diagnosed in older children (> 6 years)^21^, highlighting that VEO-IBD likely involves unique perturbations that differ from IBD in older children. Consistent with their findings, we also observed a higher abundance of OTUs of the genus *Veillonella* and *Clostridium*. Interestingly, alpha diversity did not differ between VEO-IBD and healthy controls, which contrasts earlier reports in VEO-IBD^21^ and pediatric IBD^69^. However, within the VEO-IBD group, we found a positive correlation between age and microbial diversity– a pattern not observed in healthy controls. This suggests that microbial diversification in VEO-IBD may be delayed, potentially due to early life dysbiosis combined with prolonged exposure to chronic intestinal inflammation and/or medications prescribed to control the symptoms of IBD, disrupting normal microbiome development.

We observed a lower abundance of *N*-acetyl-Met alongside a higher abundance of 5’-methylthioadenosine (MTA) in VEO-IBD, indicating a potential disruption in the methionine cycle. Additionally, microbially derived metabolites of aminosalicylate drugs^49^ were among the key discriminant features of the VEO-IBD group. Our multi-omics analysis also identified several unknown metabolic features that strongly correlated with microbial genera either enriched or depleted in VEO-IBD. Notably, using tissueMASST, we found that one of these metabolites – enriched in VEO-IBD – was also more frequently detected in the fecal samples from individuals with IBD compared to healthy subjects when searching its spectra against publicly available metabolomics datasets within the GNPS/MassIVE ecosystem. To explore the microbial origin of this feature, we searched microbeMASST and identified matches across multiple bacterial phyla, including *Veillonella* and *Bacteroides*, both of which were more abundant in VEO-IBD. Based on SIRIUS predictions and manual inspection of the MS/MS spectra, this metabolite is likely a modified tripeptide. However, further investigation is needed to confirm its structure and understand its putative role in the pathophysiology of VEO-IBD.

We note limitations of this study. First, metabolite annotations are based on MS/MS spectral matches with the GNPS spectral libraries, representing a level 2-3 annotation according to the Metabolomics Standard Initiative (MSI)^70^. Generally, MS/MS data cannot differentiate between stereoisomers, such as acetaminophen and 2-acetamidophenol. The poor alignment between acetaminophen detection and reported use (38%) may be explained by the parameters used for the FBMN job, which required at least 5 matching fragments for annotation, or inaccuracies in self-reported use of acetaminophen. Factors that can influence the gut microbiome composition and functionality, such as antibiotic exposure, probiotics intake, and dietary modifications were not reported in detail. Finally, the study has a relatively small sample size, and findings should be validated in a larger, independent cohort.

In conclusion, this is the first study investigating changes in the fecal microbiome and metabolome in children diagnosed with VEO-IBD compared to age- and sex-matched healthy controls. We identified distinct fecal microbial and metabolic signatures in VEO-IBD. Notably, we observed a marked depletion of short-chain *N*-acyl lipids, along with elevated levels of several di- and tripeptides and oxo BAs. Our integrative multi-omic analysis highlights a strong connection between gut microbial dysbiosis and metabolic alterations in VEO-IBD. Importantly, several of these features appear to be unique to VEO-IBD, distinguishing it from IBD diagnosed in older children or adults. Manipulation of gut microbiota composition and function via pre/probiotics, fecal microbiota transplantation, or antibiotics may provide an opportunity to target processes that contribute to disease development and/or severity in children with VEO-IBD^71,72^. In general, these microbiome-directed therapies have been shown to be well-tolerated with few side effects, making them attractive treatment options in the management of VEO-IBD^71^. Overall, the findings enhance our understanding of the distinct disease biology of VEO-IBD and provide a foundation for the development of more targeted diagnostic and therapeutic strategies.

## Acknowledgement

The authors would like to acknowledge support from Laura Stewart, Rendie McHenry, Dan Payne and Jim Chappell (VUMC) with healthy donor sample collection.

## Author contributions

J.M. conceptualized the study. M.N., J.A.C., and N.H. recruited patients and oversaw sample collection. Samples were processed by L.S.Z., M.C.C., and C.E. K.E.K., J.Z., V.C. performed sample extraction and LC-MS/MS analysis. K.E.K. conducted untargeted metabolomics analysis. K.E.K. and S.Z. conducted microbiome analysis. K.E.K. drafted the manuscript. P.C.D., S.M.T., and J.M. acquired funding and supervised this project. All authors provided feedback, reviewed, and approved the manuscript.

## Data availability

Untargeted LC-MS/MS data generated in this study are publicly available at GNPS/MassIVE (https://massive.ucsd.edu/) under the accession code MSV000097610. The associated FBMN job is publicly available at GNPS2: https://gnps2.org/status?task=8f1989e83449460a9a5748cd8f32df30. Additional information and processing pipelines for the microbiome data are available in Qiita: VEOIBD - ID 15748.

## Code availability

The code used for data analysis and to generate the figures can be found in the GitHub repository: https://github.com/kinekvitne/manuscript_VEO-IBD

## REFERENCES

1. Burisch J., Jess T., Martinato M., Lakatos PL., ECCO -EpiCom. The burden of inflammatory bowel disease in Europe. J Crohns Colitis 2013;7(4):322–37.

2. Muise AM., Snapper SB., Kugathasan S. The age of gene discovery in very early onset inflammatory bowel disease. Gastroenterology 2012;143(2):285–8.

3. Uhlig HH., Schwerd T., Koletzko S., Shah N., Kammermeier J., Elkadri A., et al. The diagnostic approach to monogenic very early onset inflammatory bowel disease. Gastroenterology 2014;147(5):990–1007.e3.

4. Cucinotta U., Arrigo S., Dipasquale V., Gramaglia SMC., Laganà F., Romano C., et al. Clinical course of very early-onset inflammatory bowel disease. J Pediatr Gastroenterol Nutr 2023;76(5):590–5.

5. Kelsen JR., Sullivan KE., Rabizadeh S., Singh N., Snapper S., Elkadri A., et al. NASPGHAN position paper on the evaluation and management for patients with very early-onset inflammatory bowel disease (VEO-IBD). J Pediatr Gastroenterol Nutr 2019;70(3):389–403.

6. Kammermeier J., Dziubak R., Pescarin M., Drury S., Godwin H., Reeve K., et al. Phenotypic and genotypic characterisation of inflammatory bowel disease presenting before the age of 2 years. J Crohns Colitis 2017;11(1):60–9.

7. Ouahed J., Spencer E., Kotlarz D., Shouval DS., Kowalik M., Peng K., et al. Very early onset inflammatory bowel disease: A clinical approach with a focus on the role of genetics and underlying immune deficiencies. Inflamm Bowel Dis 2020;26(6):820–42.

8. Oliva-Hemker M., Hutfless S., Al Kazzi ES., Lerer T., Mack D., LeLeiko N., et al. Clinical presentation and five-year therapeutic management of very early-onset inflammatory bowel disease in a large north American cohort. J Pediatr 2015;167(3):527–32.e1–3.

9. Rosen MJ., Dhawan A., Saeed SA. Inflammatory bowel disease in children and adolescents. JAMA Pediatr 2015;169(11):1053–60.

10. Nambu R., Warner N., Mulder DJ., Kotlarz D., McGovern DPB., Cho J., et al. A systematic review of monogenic inflammatory bowel disease. Clin Gastroenterol Hepatol 2022;20(4):e653–63.

11. Sýkora J., Pomahačová R., Kreslová M., Cvalínová D., Štych P., Schwarz J. Current global trends in the incidence of pediatric-onset inflammatory bowel disease. World J Gastroenterol 2018;24(25):2741–63.

12. Benchimol EI., Mack DR., Nguyen GC., Snapper SB., Li W., Mojaverian N., et al. Incidence, outcomes, and health services burden of very early onset inflammatory bowel disease. Gastroenterology 2014;147(4):803–13.e7; quiz e14–5.

13. Charbit-Henrion F., Parlato M., Hanein S., Duclaux-Loras R., Nowak J., Begue B., et al. Diagnostic yield of next-generation sequencing in very early-onset inflammatory bowel diseases: A multicentre study. J Crohns Colitis 2018;12(9):1104–12.

14. Lloyd-Price J., Arze C., Ananthakrishnan AN., Schirmer M., Avila-Pacheco J., Poon TW., et al. Multi-omics of the gut microbial ecosystem in inflammatory bowel diseases. Nature 2019;569(7758):655–62.

15. Khor B., Gardet A., Xavier RJ. Genetics and pathogenesis of inflammatory bowel disease. Nature 2011;474(7351):307–17.

16. Cummings JH., Macfarlane GT. Role of intestinal bacteria in nutrient metabolism. JPEN J Parenter Enteral Nutr 1997;21(6):357–65.

17. Wilson ID., Nicholson JK. Gut microbiome interactions with drug metabolism, efficacy, and toxicity. Transl Res 2017;179:204–22.

18. Zheng D., Liwinski T., Elinav E. Interaction between microbiota and immunity in health and disease. Cell Res 2020;30(6):492–506.

19. Zarrinpar A., Chaix A., Xu ZZ., Chang MW., Marotz CA., Saghatelian A., et al. Antibiotic-induced microbiome depletion alters metabolic homeostasis by affecting gut signaling and colonic metabolism. Nat Commun 2018;9(1):2872.

20. Tamburini S., Shen N., Wu HC., Clemente JC. The microbiome in early life: implications for health outcomes. Nat Med 2016;22(7):713–22.

21. Conrad MA., Bittinger K., Ren Y., Kachelries K., Vales J., Li H., et al. The intestinal microbiome of inflammatory bowel disease across the pediatric age range. Gut Microbes 2024;16(1):2317932.

22. Duboc H., Rajca S., Rainteau D., Benarous D., Maubert M-A., Quervain E., et al. Connecting dysbiosis, bile-acid dysmetabolism and gut inflammation in inflammatory bowel diseases. Gut 2013;62(4):531–9.

23. Gentry EC., Collins SL., Panitchpakdi M., Belda-Ferre P., Stewart AK., Carrillo Terrazas M., et al. Reverse metabolomics for the discovery of chemical structures from humans. Nature 2024;626(7998):419–26.

24. Bjerrum JT., Wang Y., Hao F., Coskun M., Ludwig C., Günther U., et al. Metabonomics of human fecal extracts characterize ulcerative colitis, Crohn’s disease and healthy individuals. Metabolomics 2015;11(1):122–33.

25. Marchesi JR., Holmes E., Khan F., Kochhar S., Scanlan P., Shanahan F., et al. Rapid and noninvasive metabonomic characterization of inflammatory bowel disease. J Proteome Res 2007;6(2):546–51.

26. Kolho K-L., Pessia A., Jaakkola T., de Vos WM., Velagapudi V. Faecal and serum metabolomics in paediatric inflammatory bowel disease. J Crohns Colitis 2017;11(3):321–34.

27. Chambers MC., Maclean B., Burke R., Amodei D., Ruderman DL., Neumann S., et al. A cross-platform toolkit for mass spectrometry and proteomics. Nat Biotechnol 2012;30(10):918–20.

28. Schmid R., Heuckeroth S., Korf A., Smirnov A., Myers O., Dyrlund TS., et al. Integrative analysis of multimodal mass spectrometry data in MZmine 3. Nat Biotechnol 2023;41(4):447–9.

29. Nothias L-F., Petras D., Schmid R., Dührkop K., Rainer J., Sarvepalli A., et al. Feature-based molecular networking in the GNPS analysis environment. Nat Methods 2020;17(9):905–8.

30. Wang M., Carver JJ., Phelan VV., Sanchez LM., Garg N., Peng Y., et al. Sharing and community curation of mass spectrometry data with Global Natural Products Social Molecular Networking. Nat Biotechnol 2016;34(8):828–37.

31. Shannon P., Markiel A., Ozier O., Baliga NS., Wang JT., Ramage D., et al. Cytoscape: a software environment for integrated models of biomolecular interaction networks. Genome Res 2003;13(11):2498–504.

32. Dührkop K., Fleischauer M., Ludwig M., Aksenov AA., Melnik AV., Meusel M., et al. SIRIUS 4: a rapid tool for turning tandem mass spectra into metabolite structure information. Nat Methods 2019;16(4):299–302.

33. Dührkop K., Nothias L-F., Fleischauer M., Reher R., Ludwig M., Hoffmann MA., et al. Systematic classification of unknown metabolites using high-resolution fragmentation mass spectra. Nat Biotechnol 2021;39(4):462–71.

34. Wang M., Jarmusch AK., Vargas F., Aksenov AA., Gauglitz JM., Weldon K., et al. Mass spectrometry searches using MASST. Nat Biotechnol 2020;38(1):23–6.

35. Zuffa S., Schmid R., Bauermeister A., P Gomes PW., Caraballo-Rodriguez AM., El Abiead Y., et al. microbeMASST: a taxonomically informed mass spectrometry search tool for microbial metabolomics data. Nat Microbiol 2024;9(2):336–45.

36. Mildau K., van der Hooft JJJ., Flasch M., Warth B., El Abiead Y., Koellensperger G., et al. Homologue series detection and management in LC-MS data with homologueDiscoverer. Bioinformatics 2022;38(22):5139–40.

37. Mohanty I., Mannochio-Russo H., Schweer JV., El Abiead Y., Bittremieux W., Xing S., et al. The underappreciated diversity of bile acid modifications. Cell 2024;187(7):1801–18.e20.

38. Mannochio-Russo H., Charron-Lamoureux V., van Faassen M., Lamichhane S., Nunes WDG., Deleray V., et al. The microbiome diversifies N-acyl lipid pools - including short-chain fatty acid-derived compounds. bioRxivorg 2024:2024.10.31.621412.

39. Jarmusch AK., Aron AT., Petras D., Phelan VV., Bittremieux W., Acharya DD., et al. A universal language for finding mass spectrometry data patterns. bioRxiv 2022:2022.08.06.503000. Doi: 10.1101/2022.08.06.503000.

40. Caporaso JG., Lauber CL., Walters WA., Berg-Lyons D., Huntley J., Fierer N., et al. Ultra-high-throughput microbial community analysis on the Illumina HiSeq and MiSeq platforms. ISME J 2012;6(8):1621–4.

41. Gonzalez A., Navas-Molina JA., Kosciolek T., McDonald D., Vázquez-Baeza Y., Ackermann G., et al. Qiita: rapid, web-enabled microbiome meta-analysis. Nat Methods 2018;15(10):796–8.

42. McDonald D., Price MN., Goodrich J., Nawrocki EP., DeSantis TZ., Probst A., et al. An improved Greengenes taxonomy with explicit ranks for ecological and evolutionary analyses of bacteria and archaea. ISME J 2012;6(3):610–8.

43. Bolyen E., Rideout JR., Dillon MR., Bokulich NA., Abnet CC., Al-Ghalith GA., et al. Reproducible, interactive, scalable and extensible microbiome data science using QIIME 2. Nat Biotechnol 2019;37(8):852–7.

44. McMurdie PJ., Holmes S. phyloseq: an R package for reproducible interactive analysis and graphics of microbiome census data. PLoS One 2013;8(4):e61217.

45. Anderson MJ. Permutational multivariate analysis of variance (PERMANOVA). Wiley StatsRef: Statistics Reference Online 2017:1–15. Doi: 10.1002/9781118445112.stat07841.

46. Fernandes AD., Reid JN., Macklaim JM., McMurrough TA., Edgell DR., Gloor GB. Unifying the analysis of high-throughput sequencing datasets: characterizing RNA-seq, 16S rRNA gene sequencing and selective growth experiments by compositional data analysis. Microbiome 2014;2(1):15.

47. Singh A., Shannon CP., Gautier B., Rohart F., Vacher M., Tebbutt SJ., et al. DIABLO: an integrative approach for identifying key molecular drivers from multi-omics assays. Bioinformatics 2019;35(17):3055–62.

48. Zhao HN., Kvitne KE., Brungs C., Mohan S., Charron-Lamoureux V., Bittremieux W., et al. Empirically establishing drug exposure records directly from untargeted metabolomics data. bioRxiv 2024:2024.10.07.617109. Doi: 10.1101/2024.10.07.617109.

49. Mehta RS., Mayers JR., Zhang Y., Bhosle A., Glasser NR., Nguyen LH., et al. Gut microbial metabolism of 5-ASA diminishes its clinical efficacy in inflammatory bowel disease. Nat Med 2023;29(3):700–9.

50. Uhlig HH. Monogenic diseases associated with intestinal inflammation: implications for the understanding of inflammatory bowel disease. Gut 2013;62(12):1795–805.

51. Bushman FD., Conrad M., Ren Y., Zhao C., Gu C., Petucci C., et al. Multi-omic analysis of the interaction between Clostridioides difficile infection and pediatric inflammatory bowel disease. Cell Host Microbe 2020;28(3):422–33.e7.

52. Bosch S., Struys EA., van Gaal N., Bakkali A., Jansen EW., Diederen K., et al. Fecal amino acid analysis can discriminate DE Novo treatment-naïve pediatric inflammatory bowel disease from controls. J Pediatr Gastroenterol Nutr 2018;66(5):773–8.

53. Ning L., Zhou Y-L., Sun H., Zhang Y., Shen C., Wang Z., et al. Microbiome and metabolome features in inflammatory bowel disease via multi-omics integration analyses across cohorts. Nat Commun 2023;14(1):7135.

54. Franzosa EA., Sirota-Madi A., Avila-Pacheco J., Fornelos N., Haiser HJ., Reinker S., et al. Gut microbiome structure and metabolic activity in inflammatory bowel disease. Nat Microbiol 2019;4(2):293–305.

55. Chang F-Y., Siuti P., Laurent S., Williams T., Glassey E., Sailer AW., et al. Gut-inhabiting Clostridia build human GPCR ligands by conjugating neurotransmitters with diet- and human-derived fatty acids. Nat Microbiol 2021;6(6):792–805.

56. Louis P., Flint HJ. Formation of propionate and butyrate by the human colonic microbiota. Environ Microbiol 2017;19(1):29–41.

57. Mills RH., Dulai PS., Vázquez-Baeza Y., Sauceda C., Daniel N., Gerner RR., et al. Multi-omics analyses of the ulcerative colitis gut microbiome link Bacteroides vulgatus proteases with disease severity. Nat Microbiol 2022;7(2):262–76.

58. Mohanty I., Allaband C., Mannochio-Russo H., El Abiead Y., Hagey LR., Knight R., et al. The changing metabolic landscape of bile acids - keys to metabolism and immune regulation. Nat Rev Gastroenterol Hepatol 2024;21(7):493–516.

59. Quinn RA., Melnik AV., Vrbanac A., Fu T., Patras KA., Christy MP., et al. Global chemical effects of the microbiome include new bile-acid conjugations. Nature 2020;579(7797):123–9.

60. Song Z., Cai Y., Lao X., Wang X., Lin X., Cui Y., et al. Taxonomic profiling and populational patterns of bacterial bile salt hydrolase (BSH) genes based on worldwide human gut microbiome. Microbiome 2019;7(1):9.

61. Guzior DV., Okros M., Shivel M., Armwald B., Bridges C., Fu Y., et al. Bile salt hydrolase acyltransferase activity expands bile acid diversity. Nature 2024;626(8000):852–8.

62. Rimal B., Collins SL., Tanes CE., Rocha ER., Granda MA., Solanki S., et al. Bile salt hydrolase catalyses formation of amine-conjugated bile acids. Nature 2024;626(8000):859–63.

63. Quinn R., Fu Y., Guzior D., Okros M., Bridges C., Rosset S., et al. Balance between bile acid conjugation and hydrolysis activity can alter outcomes of gut inflammation. Research Square 2024. Doi: 10.21203/rs.3.rs-5005563/v1.

64. Alimov I., Menon S., Cochran N., Maher R., Wang Q., Alford J., et al. Bile acid analogues are activators of pyrin inflammasome. J Biol Chem 2019;294(10):3359–66.

65. Ejderhamn J., Rafter JJ., Strandvik B. Faecal bile acid excretion in children with inflammatory bowel disease. Gut 1991;32(11):1346–51.

66. Fitzgerald RS., Sanderson IR., Claesson MJ. Paediatric inflammatory bowel disease and its relationship with the microbiome. Microb Ecol 2021;82(4):833–44.

67. Holmberg SM., Feeney RH., Prasoodanan P K V., Puértolas-Balint F., Singh DK., Wongkuna S., et al. The gut commensal Blautia maintains colonic mucus function under low-fiber consumption through secretion of short-chain fatty acids. Nat Commun 2024;15(1):3502.

68. Hidalgo-Cantabrana C., Delgado S., Ruiz L., Ruas-Madiedo P., Sánchez B., Margolles A. Bifidobacteria and their health-promoting effects. Microbiol Spectr 2017;5(3). Doi: 10.1128/microbiolspec.BAD-0010-2016.

69. Kowalska-Duplaga K., Gosiewski T., Kapusta P., Sroka-Oleksiak A., Wędrychowicz A., Pieczarkowski S., et al. Differences in the intestinal microbiome of healthy children and patients with newly diagnosed Crohn’s disease. Sci Rep 2019;9(1):18880.

70. Sumner LW., Amberg A., Barrett D., Beale MH., Beger R., Daykin CA., et al. Proposed minimum reporting standards for chemical analysis Chemical Analysis Working Group (CAWG) Metabolomics Standards Initiative (MSI): Chemical Analysis Working Group (CAWG) Metabolomics Standards Initiative (MSI). Metabolomics 2007;3(3):211–21.

71. Uhlig HH., Powrie F. Translating immunology into therapeutic concepts for inflammatory bowel disease. Annu Rev Immunol 2018;36(1):755–81.

72. Sullivan KE., Conrad M., Kelsen JR. Very early-onset inflammatory bowel disease: an integrated approach. Curr Opin Allergy Clin Immunol 2018;18(6):459–69.

