## Supplementary Figures for "Fecal Microbial and Metabolic Signatures in VEO-IBD: Implications for Unique Pathophysiology"

**Affiliations:**
[1] Skaggs School of Pharmacy and Pharmaceutical Sciences, University of California San Diego, La Jolla, CA, USA
[2] Collaborative Mass Spectrometry Innovation Center, Skaggs School of Pharmacy and Pharmaceutical Sciences, University of California San Diego, La Jolla, CA, USA
[3] Center for Microbiome Innovation, University of California San Diego, La Jolla, CA, USA
[4] Department of Obstetrics Gynecology and Reproductive Sciences, University of California San Diego, La Jolla, CA, USA
[5] Department of Pediatrics, Division of Gastroenterology, Hepatology, and Nutrition, Vanderbilt University Medical Center, Nashville, Tennessee, USA
[6] Department of Pathology, Microbiology, and Immunology, Vanderbilt University Medical Center, Nashville, Tennessee, USA
[7] Department of Pediatrics, Division of Hematology and Oncology, Vanderbilt University Medical Center, Nashville, Tennessee, USA
[8] Department of Pediatrics, Division of Infectious Diseases, Vanderbilt University Medical Center, Nashville, Tennessee, USA

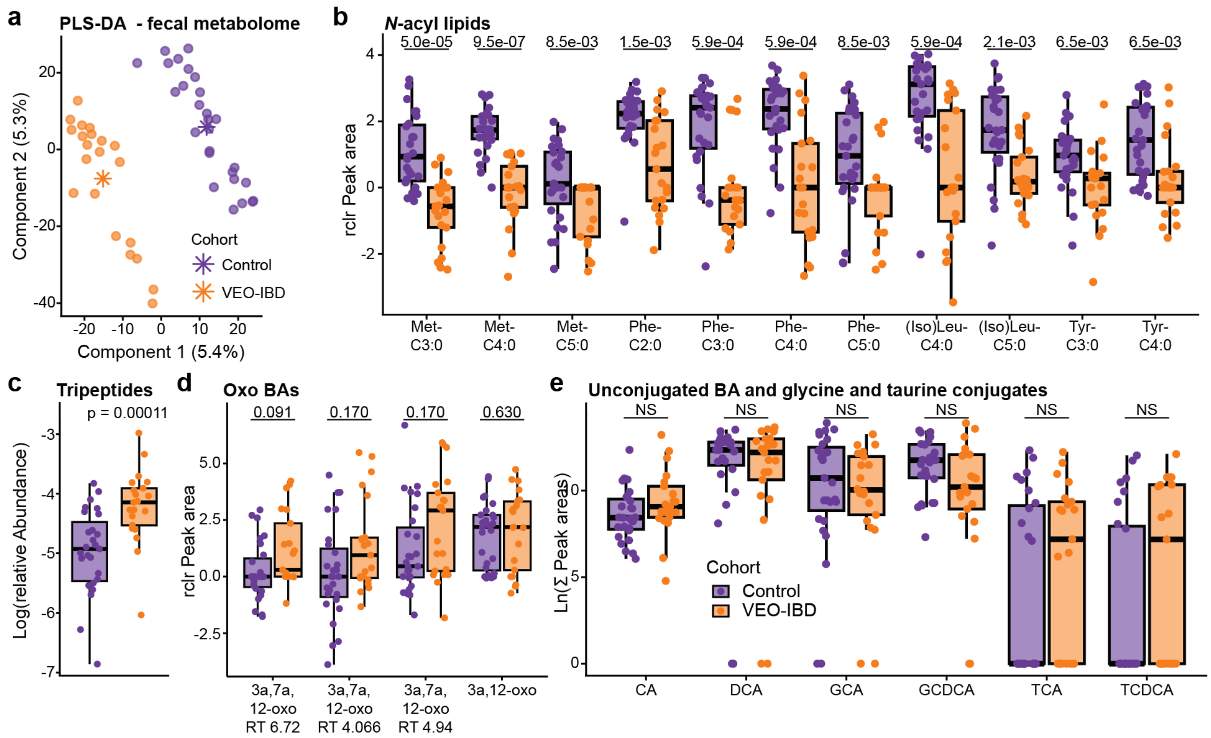


**SI Figure 1. PLS-DA of the fecal metabolome and group-wise comparisons of discriminant features and metabolite classes between VEO-IBD and healthy controls.** (a) Supervised PLS-DA of the fecal metabolome (classification error rate (CER) = 0.21). (b) Boxplot of putatively annotated short-chain *N*-acyl lipids. (c) Boxplot of the relative abundance of tripeptides with a VIP score > 1 from the PLS-DA model. Both annotated and SIRIUS predicted (probability scores > 0.7) tripeptides are included in the plot. Relative abundance was calculated by dividing the sum of peak areas for relevant tripeptides by the total extracted peak areas. (d) Boxplot of putatively annotated oxo BAs. Three of them are isomers with different retention time (RT). (e) Boxplot of MS/MS spectral matches to putative cholic acid (CA), deoxycholic acid (DCA), glycocholic acid (GCA), glycochenodeoxycholic acid (GCDCA), taurocholic acid (TCA), and taurochenodeoxycholic acid (TCDCA) (all p > 0.05). Significance between VEO-IBD and control was tested with Wilcoxon rank-sum test. P-values in (b), (d), and (e) were adjusted for multiple comparisons using the Benjamini-Hochberg correction. All boxplots show the first (lower), median, and third (upper) quartiles, with whiskers 1.5 times the interquartile range.

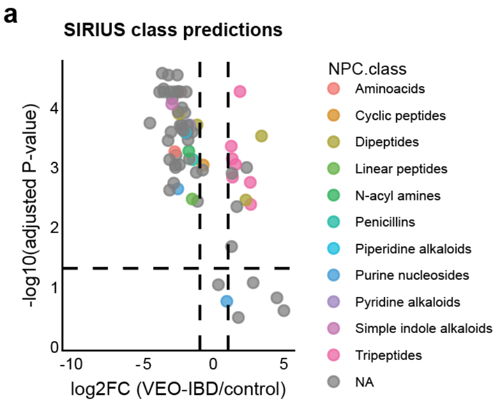


**SI Figure 2. Unannotated discriminant features by SIRIUS class prediction** (a) Volcano plot of unannotated molecules with a VIP > 2.5 colored based on SIRIUS predicted NPC class (probability > 0.7). The dotted lines show the significance threshold (p < 0.05 and absolute log2 fold change >2). P-values were adjusted for multiple comparisons using the Benjamini-Hochberg correction.

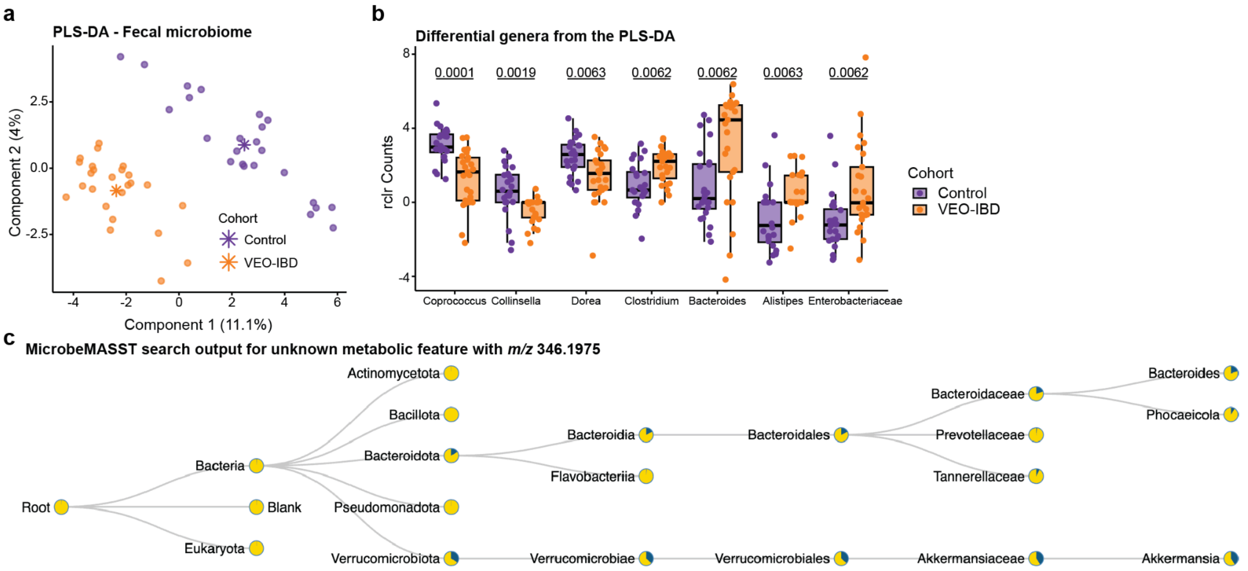


**SI Figure 3. PLS-DA of the fecal microbiome and group-wise comparisons of differential genera between VEO-IBD and healthy controls.** (a) Supervised PLS-DA of the fecal microbiome (classification error rate (CER) = 0.11). (b) Boxplot of selected discriminant genera from the PLS-DA model. All boxplots show the first (lower), median, and third (upper) quartiles, with whiskers 1.5 times the interquartile range. P-values were adjusted for multiple comparisons using the Benjamini-Hochberg correction. (c) The microbeMASST search output for an unannotated metabolic feature (*m/z* 346.1975) enriched in VEO-IBD. Pie charts show the proportion of spectral matches found in deposited datasets, with blue representing a match and yellow representing a non-match.
